# FLIMPA: A versatile software for Fluorescence Lifetime Imaging Microscopy Phasor Analysis

**DOI:** 10.1101/2024.09.13.612802

**Authors:** Sofia Kapsiani, Nino F. Läubli, Edward N. Ward, Mona Shehata, Clemens F. Kaminski, Gabriele S. Kaminski Schierle

## Abstract

Fluorescence lifetime imaging microscopy (FLIM) is an advanced microscopy technique capable of providing a deeper understanding of the molecular environment of a fluorophore. While FLIM data were traditionally analysed through the exponential fitting of the fluorophores’ emission decays, the use of phasor plots is increasingly becoming the preferred standard. This is due to their ability to visualise the distribution of fluorescent lifetimes within a sample, offering insights into molecular interactions in the sample without the need for model assumptions regarding the exponential decay behaviour of the fluorophores. However, so far most researchers have had to rely on commercial phasor plot software packages, which are closed-source and rely on proprietary data formats. In this paper, we introduce FLIMPA, an opensource, stand-alone software for phasor plot analysis that provides many of the features found in commercial software, and more. FLIMPA is fully developed in Python and offers advanced tools for data analysis and visualisation. It enhances FLIM data comparison by integrating phasor points from multiple trials and experimental conditions into a single plot, while also providing the possibility to explore detailed, localised insights within individual samples. We apply FLIMPA to introduce a cell-based assay for the quantification of microtubule depolymerisation, measured through fluorescence lifetime changes of SiR-tubulin, in response to various concentrations of Nocodazole, a microtubule depolymerising drug relevant to anti-cancer treatment.

## 1 Introduction

The fluorescence lifetime of a fluorophore is the average time a fluorophore remains in its excited state before emitting a photon and returning to the ground state [1]. Variations in fluorescence lifetime can be leveraged to gain insights into the molecular environment of the fluorophores, including changes in temperature, pH, and protein-protein interactions as measured by Förster Resonance Energy Transfer (FRET), among other factors [2, 3]. Fluorescence lifetimes are typically measured by fluorescence lifetime imaging microscopy (FLIM), a powerful optical technique that provides the spatial mapping of the fluorophores’ fluorescence lifetime within a sample [4]. FLIM is more robust than conventional fluorescence intensity techniques as it is less susceptible to experimental fluctuations, such as fluctuations in fluorophore concentration, laser intensity, and photobleaching [3, 5].

Over many years, FLIM has been applied to study a multitude of biological processes. For example, FLIM-based intracellular biosensors have been used to measure changes in amyloid aggregation state [6, 7], calcium concentration [8, 9, 10], NAD(P)H levels [11, 12], ATP cleavage [13], hydrogen peroxide levels [14, 15], redox changes in the endoplasmic reticulum [16], chromatin compaction states [4] and more. FLIM data acquisition can be performed both in the time and frequency domains. The predominate technique is time-correlated single-photon counting (TCSPC)-FLIM, which features a high photon detection efficiency and smallest temporal resolution [5, 17, 18, 19]. Similar to data acquisition, FLIM data can also be analysed either in the time or frequency domain. In the time domain, the fluorescence lifetime parameters can be extracted using curvefitting techniques through open-source software such as FLIMfit [20], FLIMJ [21], and FLIMView [22]. However, these curve-fitting techniques are computationally expensive, require user-defined input parameters and knowledge of fluorescence decay characteristics, and thus depend on high level of expertise for data quantification and interpretation [5]. An alternative method to compute the fluorescence lifetime values is via phasor plot analysis, which involves the transformation of the data from the time to the frequency domain, using Fourier transformation.

In contrast to traditional fluorescence lifetime fitting, the phasor approach does not require model fitting and, it is therefore, computationally inexpensive. Furthermore, no assumptions need to be made about the imaged fluorophore and the form of its fluorescence lifetime decay [23]. Each pixel in the image is mapped onto the two-dimensional phasor space, enabling the visualisation of the distribution of the fluorescence lifetimes in the sample and the detection of small but significant differences between populations [19, 2]. This visualisation facilitates a deeper understanding of the identities and behaviours of fluorophores present in the sample [24]. However, currently, most software implementations available for phasor analysis, such as SPCImage [25], Leica LAS X [26], FLIM STUDIO [27] and VistaVision [23] are part of commercial microscopy systems, limiting adjustability and flexibility and restricting access to their proprietary file formats [28]. Recently, the open-source Python-based software, FLUTE [28], a Napari-Live-FLIM plugin [29], and Phasor identifier [30] have been published, highlighting a demenad for open-source phasor plot analysis software. Unfortunately, the latter offer only limited options for data analysis and usability compared with their commercial counterparts or they are not stand-alone applications that can be used without programming expertise. To address these limitations, we introduce FLIMPA, an open-source, user-friendly, and stand-alone software package for phasor plot analysis of TCSPC-FLIM data. FLIMPA features advanced options for data analysis, visualisation, and interpretation, which include the ability to highlight specific regions of interest (ROI) within individual images, display gallery plots that show multiple images in a single view, tools for manual mask addition, and the generation of violin plots to visualise the data distributions for different experimental conditions or phenotypes. The results are displayed across several tabs, in a similar format to what is used in the widely popular FLIMfit [20] software, ensuring accessibility and easy transition for FLIM researchers [20]. Further, to ensure FLIMPA’s usability across different laboratories and imaging setups, a number of file formats are supported, currently, .sdt (Becker & Hickl), .ptu (PicoQuant), and .tif.

To highlight FLIMPA’s capabilities, we have performed a case study of how to quantify the state of microtubule depolymerisation upon drug exposure via a small molecule dye, SiR-tubulin. SiR-tubulin is non-toxic and selectively stains microtubules in live cells without requiring genetic modification. Microtubules are key components of the cell’s cytoskeleton, crucial for transport and sensing within the cell. They also form the mitotic spindle and thus facilitate cell division [31, 32]. Microtubules are therefore important targets of anti-cancer drugs as disruption of microtubule dynamics can halt cell division, leading to tumour death [33, 34], as illustrated by Supplementary Figure 1.

In this study, Nocodazole, a microtubule depolymerising agent, was utilised to induce microtubule disassembly in live COS-7 cells. Nocodazole has been shown to bind to unpolymerised tubulin within the cells, thereby disrupting microtubule assembly [35]. Changes in the fluorescence lifetime of SiR-tubulin following the addition of Nocodazole have been used to quantify the degree of microtubule depolymerisation [35]. As a Rhodamine-based dye, SiR-tubulin can undergo self-quenching when dye molecules are in close proximity, leading to a change in the fluorescence lifetime [36]. Consequently, if the microtubules remain intact, SiR-tubulin strongly self-quenches due to the proximity of neighbouring dye molecules when they are attached to the microtubules. On the other hand, Nocodazole leads to a dispersion of the tubulin subunits, and thus a reduction in the self-quenching between the SiR-tubulin molecules with a concomitant increase in fluorescence lifetime. Similar observations have been made for amyloid proteins imaged using a rhodmanine based dye which self quenches upon amyloid aggregation [6]. This is the first application of FLIM for building a cell-based assay for measuring drug-induced microtubule destabilisation.

## 2 Results

FLIMPA is designed for phasor plot analysis of raw TCSPC-FLIM data. It is open-source and can be installed on a Windows computer using the .exe file available on its GitHub repository. An overview of the graphical user interface (GUI) is provided in Figure 1. For proof-of-principle analysis, we used a specimen of Convallaria Rhizome and compared FLIMPA’s output with the fluorescence lifetime map obtained using FLIMfit [20], achieving similar results (Supplementary Figure 2). The colour bars for the fluorescence lifetime maps are in nanoseconds (ns), while the colour bars of the intensity images represent the number of photons per pixel.

**Figure 1:**
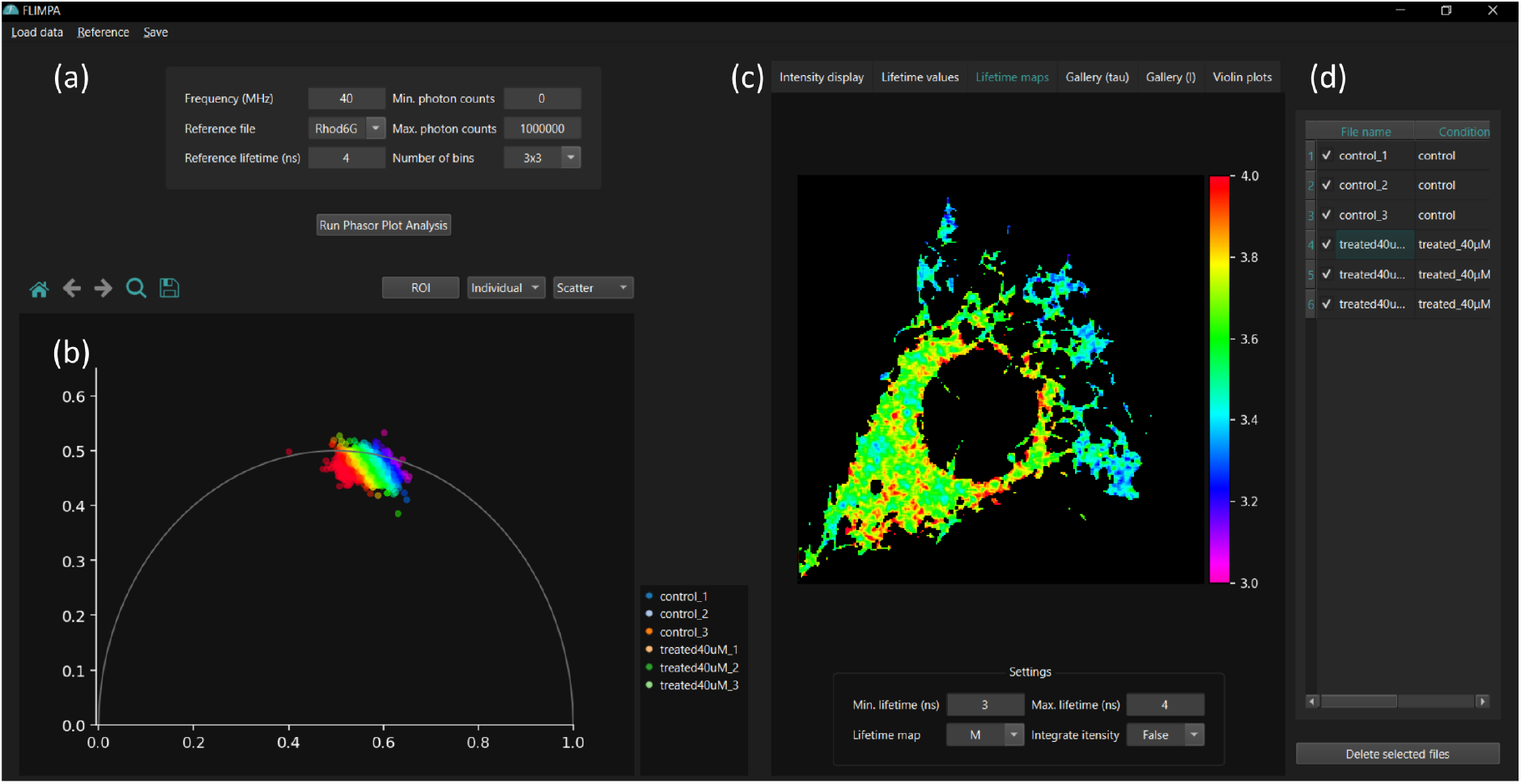
The FLIMPA software provides an easy to use user interface, where: (a) displays the area for setting imaging and analysis parameters, including laser repetition rate/frequency, reference file, and photon count thresholds; (b) shows the phasor plot visualisation, with the x− and y−axis representing the G− and S− phasor coordinates (please refer to Phasor Plot Theory section in Methods); (c) depicts the visualisation options post-analysis, including the fluorescence lifetime and intensity maps; (d) stores the imported file names and their corresponding experimental conditions, if provided.

As shown in Figure 1b, the phasor plots are displayed on the left-hand side of the GUI, while additional visualisations are presented across different tabs on the right-hand side (Figure 1c). An overview of the different tabs is shown in Figure 2, which include a table listing the phase, modulation, and average fluorescence lifetime values (see Phasor Plot Theory section in Methods) for each image or ROI, visualisation of fluorescence lifetime and intensity maps as well as violin plots, to compare fluorescence lifetime distributions between different data sets.

FLIMPA serves as a powerful tool for the analysis of bulk FLIM data, offering the capability to visualise phasor plots from single images as well as combined plots from multiple samples and datasets. This allows users to explore localised effects within individual images, such as the presence of undesired autofluorescence or local quenching related to excess in concentrations of dye molecules. It is thus easy to examine how the fluorescence lifetime varies across different experimental conditions. The phasor plot analysis requires a reference file of a dye with a known fluorescence lifetime to account for instrumental errors. Typically, dyes exhibiting mono-exponential decay kinetics, such as Rhodamine 6G (fluorescence lifetime of 4 ns in water [37]) or Erythrosin B (fluorescence lifetime of 0.089 ns in water [38]), are used for reference measurements and system calibration. Prior to the analysis, users can also mask the background of samples based on the number of photons per pixel by setting minimum and maximum thresholds, as shown in Figure 3(b), thus excluding, for example, autofluorescence which is usually present at low photon counts. FLIMPA also accepts manual masks to define multiple regions of interest (ROI) (Figure 3c).

**Figure 2:**
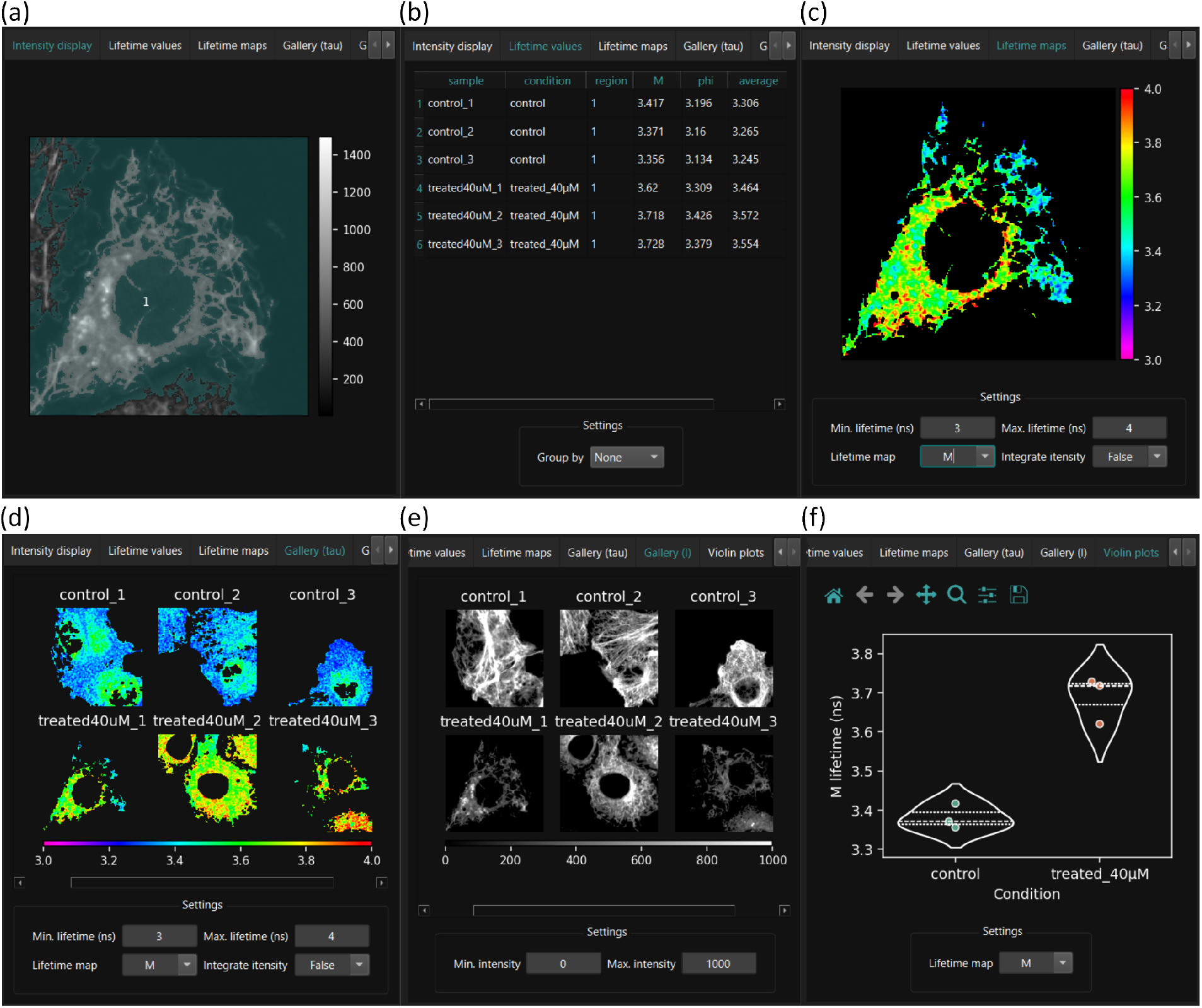
Different tabs of the FLIMPA software permit robust and versatile data analysis. (a) “Intensity display” depicting the input data with intensity masking, if masking has been applied. (b) “Lifetime values” is the output table displaying the mean modulation, phase and average fluorescence lifetime per image. Additionally, the mean fluorescence lifetime, per ROI (if different ROIs have been selected using manual masking) or per experimental condition, can be shown. (c) “Lifetime maps” show the individual images where localised effects can be explored using the ROI selection tool (please refer to section 2.1). (d) “Gallery (tau)” is a gallery of the fluorescence lifetime maps for the different samples analysed. (e) “Gallery (I)” is a gallery of the fluorescence intensity maps. (f) “Violin plots” shows the distribution of data points for each treatment group.

**Figure 3:**
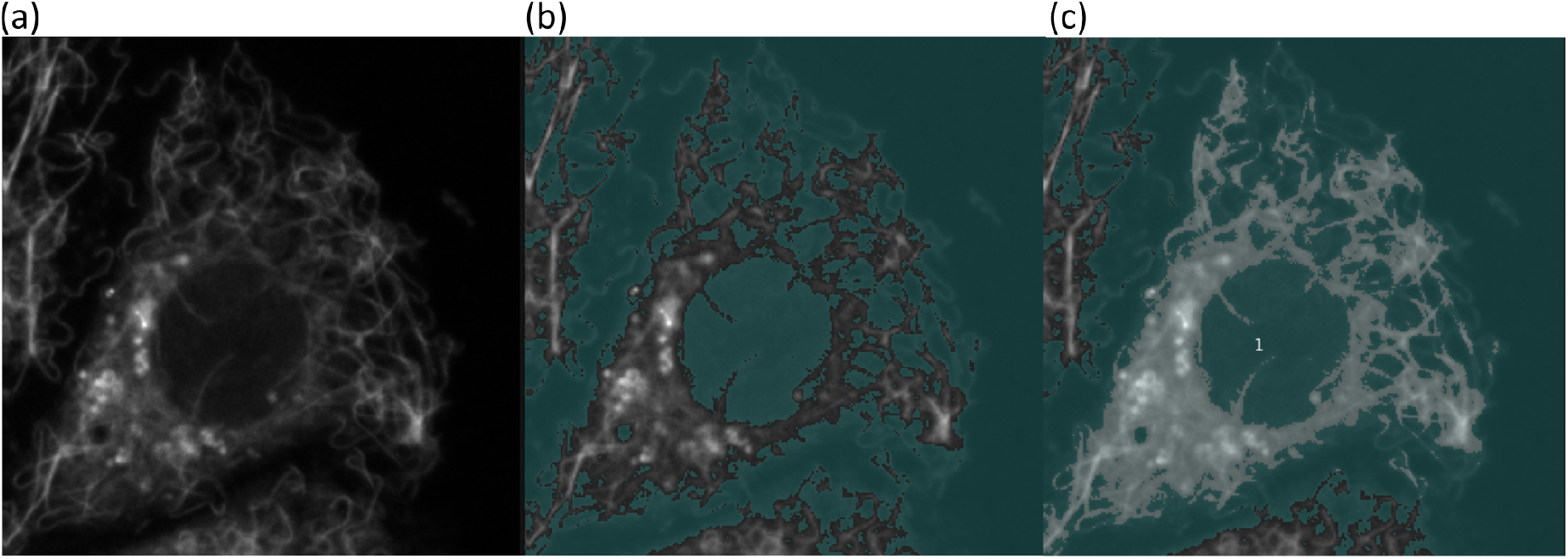
The “intensity display” tab of FLIMPA permits the selection of individual identity masks for improved data analysis. (a) Shows the raw, unmasked image of COS-7 cells displaying SiR-tubulin labelled microtubules treated with 40 *µ*M Nocodazole. (b) Displays the image with background masking applied by setting a minimum photon count threshold per pixel to reduce the level of autofluorescence background. The masked area is shown in dark blue-green. (c) Illustration of an additional, manual mask being applied. In this example, only the central cell (highlighted) has been selected for data analysis.

Upon loading the sample data into FLIMPA, users have the option to assign experimental conditions to each sample, thus allowing samples of the same set of conditions to be visualised together. FLIMPA also supports pixel binning, as commonly applied in FLIMfit [20], to aid in the analysis of data with lower signal-to-noise ratios.

### 2.1 FLIMPA enables the investigation of localised fluorescence lifetime changes within individual images

Localised effects of treatment within individual images can be studied using the fluorescence “lifetime maps” tab, where the output images are pseudo-coloured based on the fluorescence lifetime value for each image pixel. Alongside the lifetime maps, the corresponding phasor plot is displayed, where the phasor plot is plotted with colours that map to corresponding locations on the lifetime images. Furthermore, FLIMPA allows for the selection of specific ROI on the phasor plot. The pixels within the selected ROI are highlighted on the lifetime map, enabling the user to investigate localised changes in fluorescence lifetime and to correlate them with specific cellular structures or localised treatment effects. Hence, the ROI selection tool can aid in the identification of outliers. As illustated in Figure 4a, several pixels of lower fluorescence lifetimes, coloured in pink, are shown as separated from the main phasor cluster.

**Figure 4:**
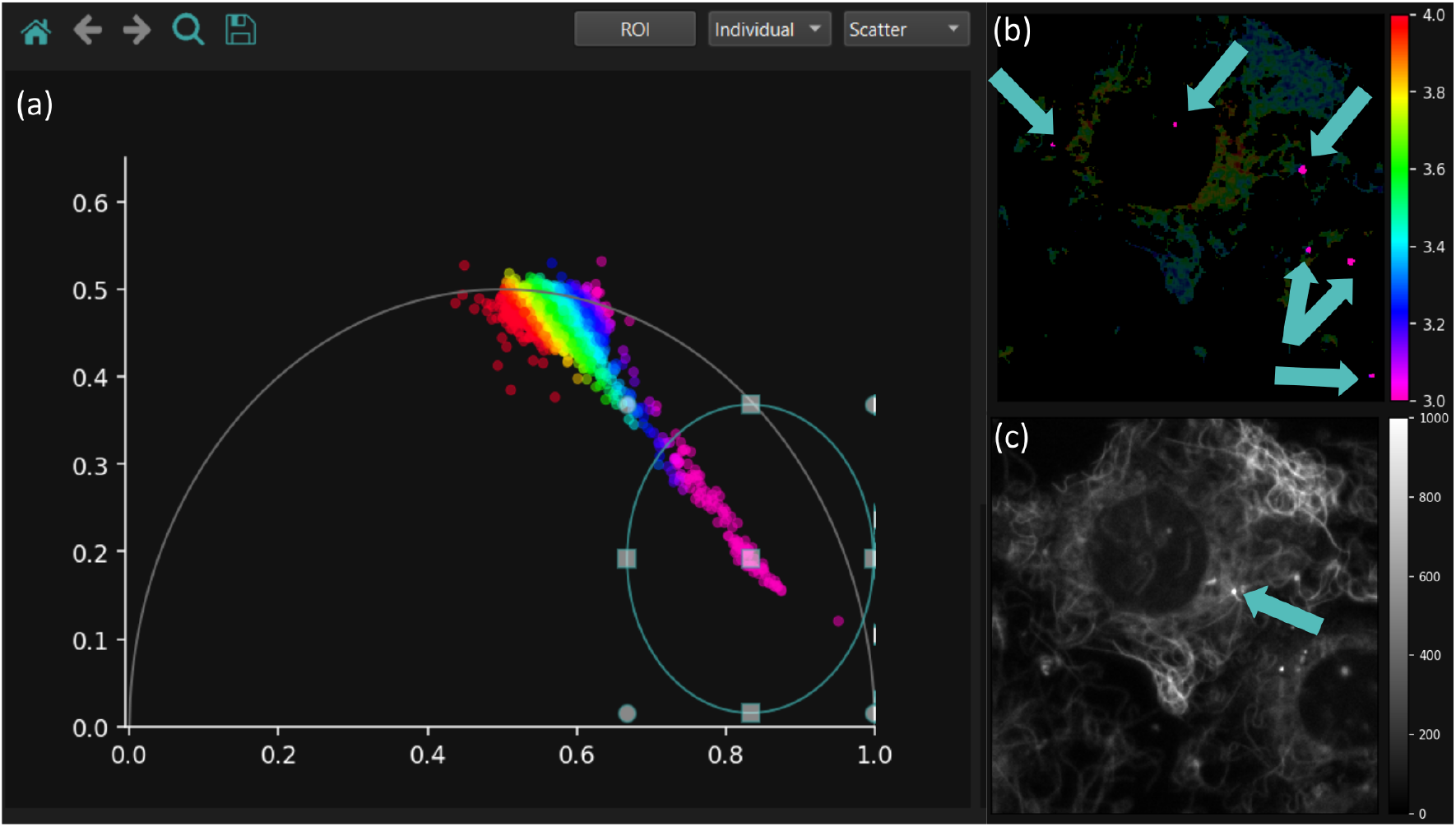
The ROI selection tool can be utilised to explore different regions within the sample to visualise localised treatment effects or to detect localised outliers. (a) Phasor points with lower fluorescence lifetimes compared to the main phasor cluster have been selected using the ROI tool. (b) The modulation lifetime map highlights pixels corresponding to the ROI-selected phasor in pink. These regions indicated by arrows likely represent SiR-tubulin aggregates and should be excluded from the data analysis. (c) The intensity image, with a blue arrow indicating the microtubule organising centre (MTOC), confirms that the phasor points with lower lifetime do not represent the cells’ MTOCs.

The individual dots observed through the phasor plot-based segmentation likely correspond to aggregated SiR-tubulin monomers that have been detached from the cells. SiR-tubulin aggregation increases self-quenching, resulting in a decrease in the fluorescence lifetime. Indeed, Pineda *et al*. (2018) [39] also reported that SiR-tubulin is prone to aggregation and recommended centrifuging the dye before cell staining to mitigate this effect. Additionally, the intensity image shown in Figure 4c is used to verify that these regions do not correspond to MTOCs, which, due to their tight arrangement, could also display a lower fluorescence lifetime.

### 2.2 FLIMPA is a powerful tool for studying global trends across different experimental conditions

To investigate how different phenotypes or experimental conditions affect a probe’s fluorescence lifetime, FLIMPA enables the visualisation of collective data from different measurements in a single phasor plot. Users can customise the colouring of the phasor points according to sample identity (Figure 5a), where the phasor cloud for each sample is visualised with a different colour. Alternatively, the points can be coloured in alignment with their experimental condition and combined into a single phasor cloud, allowing comparisons across experimental groups (Figure 5b).

**Figure 5:**
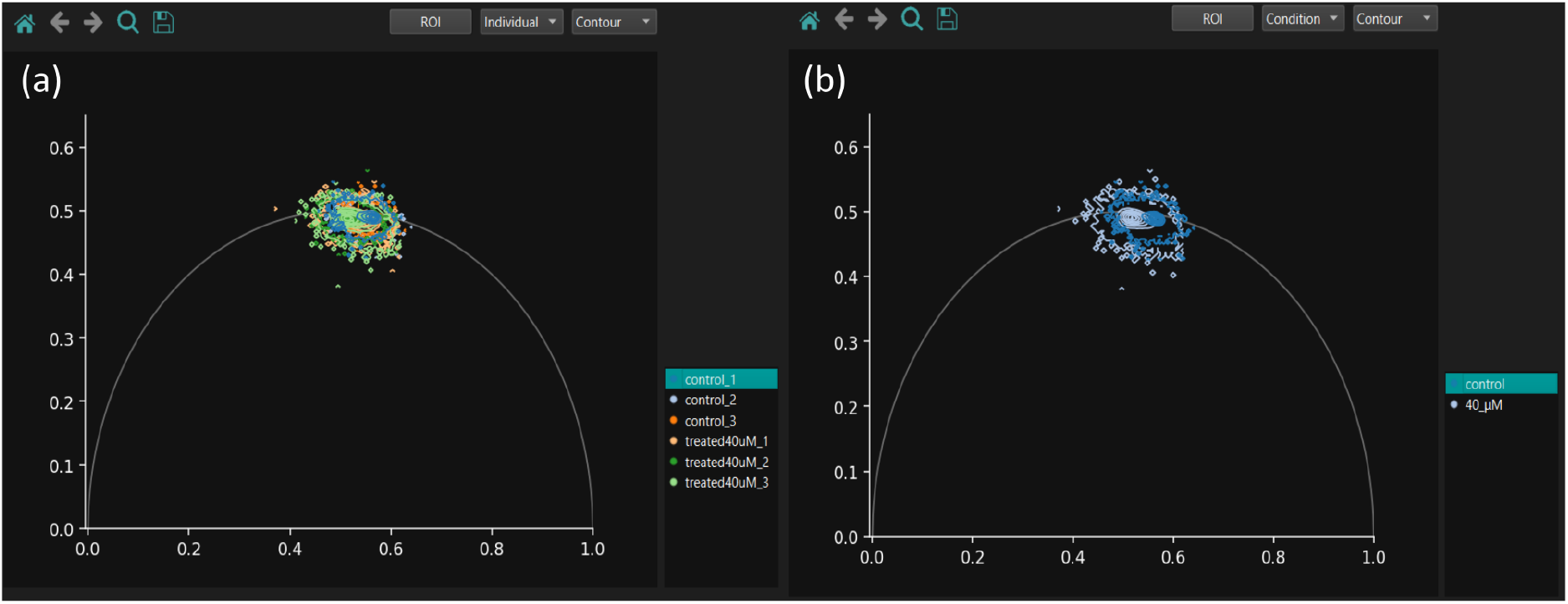
FLIMPA provides data visualisation options to assist in bulk data analysis and interpretation.(a) Phasor plot visualisation, where each contour map is coloured based on sample identity. (b) The contour maps are coloured according to the assigned experimental condition (light blue for control and darker blue for 40 *µ*M of Nocodazole).

In addition to the contour maps displayed in Figure 5, FLIMPA offers scatter plot and density-sensitive histogram visualisation options (Supplementary Figure 3). As for the fluorescence lifetime and intensity maps, all phasor plots can be exported as .png files. Further, the figures can be saved with a transparent background to aid the subsequent merging of different conditions in individual plots for comparison.

### 2.3 Phasor analysis reveals increase in SiR-tubulin fluorescence lifetime with increasing Nocodazole concentrations

Here, FLIMPA has been used to analyse how different concentrations of Nocodazole, specifically, 0 *µ*M (control), 1 *µ*M, 10 *µ*M, and 40 *µ*M, affect microtubule stability. The exported phasor plots are shown in Figures 6 (a-d).

**Figure 6:**
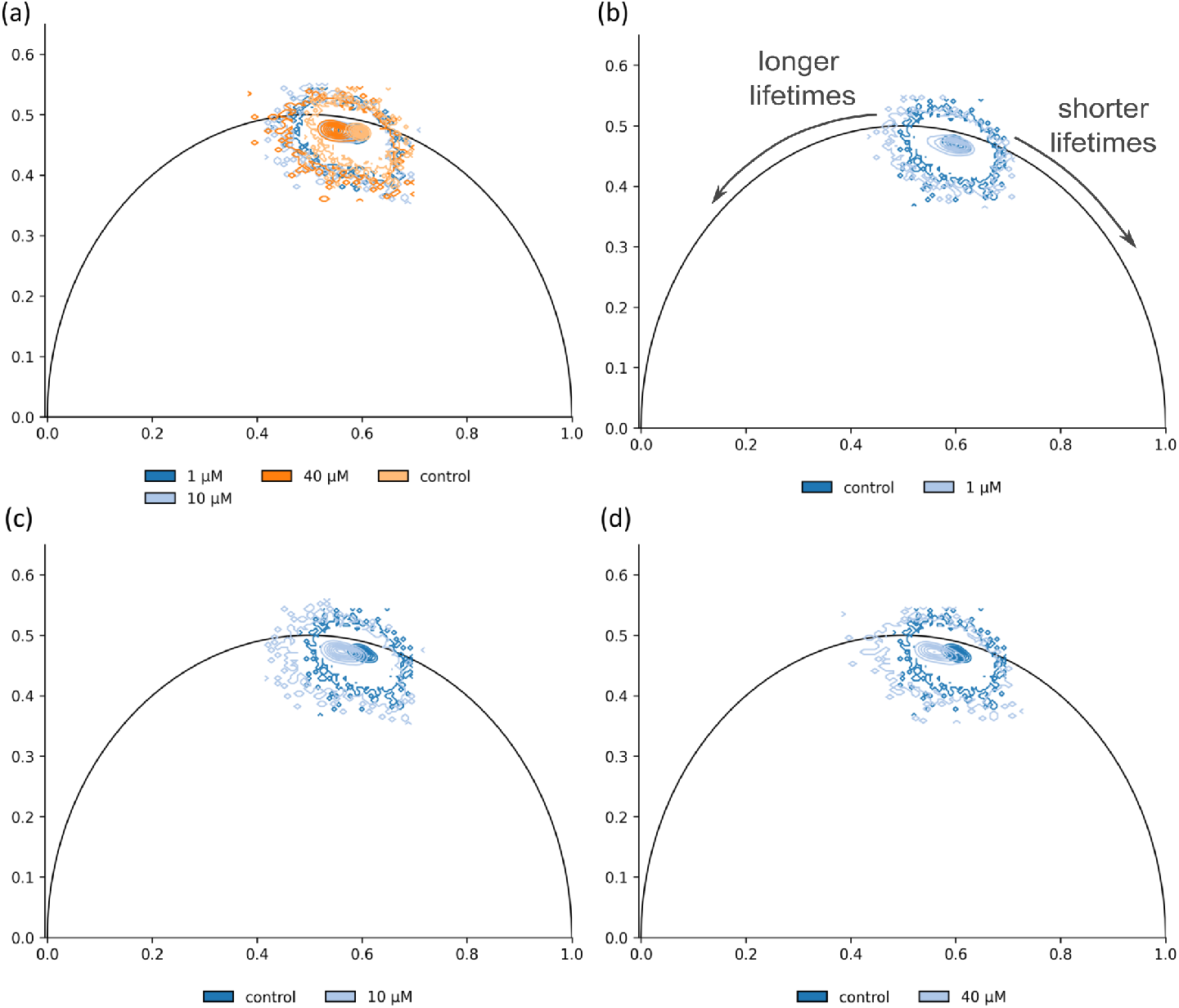
FLIMPA reveals that increasing concentration of Nocodazole significantly destabilises microtubules. Phasor plots exported from FLIMPA displaying contour maps for (a) control (light orange), 1 *µ*M Nocodazole (darker blue), 10 *µ*M Nocodazole (light blue), and 40 *µ*M Nocodazole (darker orange); (b) control (darker blue) versus 1 *µ*M Nocodazole (light blue); (c) control (darker blue) versus 10 *µ*M Nocodazole (light blue); and (d) control (darker blue) versus 40 *µ*M Nocodazole (light blue). The figure was edited in Inkscape to indicate the shift towards longer and shorter lifetimes.

Specifically, Nocodazole treatment shifted the distribution of fluorescence lifetimes towards the left-hand side of the phasor plot’s universal circle, indicating longer lifetimes compared to the control condition (see Phasor Plot Theory section for more information). This effect was more pronounced for phasor clusters corresponding to 10 *µ*M and 40 *µ*M Nocodazole treatments than for the 1 *µ*M Nocodazole, indicating that the former concentrations induced a higher degree of microtubule depolymerisation. Moreover, the phasor plots highlighted that Nocodazole treatment had a strong effect on both the phase and modulation lifetimes, as indicated by a change in both the length and the angle of the phasor cloud on the plot, respectively. Specifically, the shift in length of the phasor reflects an alteration in the modulation lifetime, while the change in angle corresponds to a change in the phase lifetime.

Further statistical analysis was performed by exporting a .csv file containing the fluorescence lifetime values for each image and applying a one-way ANOVA test with Tukey multiple comparisons in Python. We note that FLIMPA does not include a possibility for statistical evaluations. This is intentional, as the selection of appropriate tests is highly dependent on the analysed dataset and its data distribution. We recommend the use of commonly used software for these tests either commercial available or using custom Python code or R packages which are readily available. Here, we focused on the modulation lifetime for analysis, as it exhibited a higher sensitivity to Nocodazole treatment. The performed ANOVA confirmed significant differences (p-value *<* 0.05) between the control and each of the treated conditions (Figure 7a), showing a statistically significant increase in the modulation lifetime of SiR-tubulin from treatment with 1 *µ*M Nocodazole to 10 *µ*M Nocodazole. However, treating the COS-7 cells with 40 *µ*M Nocodazole did not result in a further significant increase in modulation lifetime compared to 10 *µ*M Nocodazole, indicating that 10 *µ*M Nocodazole is enough to completely destabilise the microtubules.

**Figure 7:**
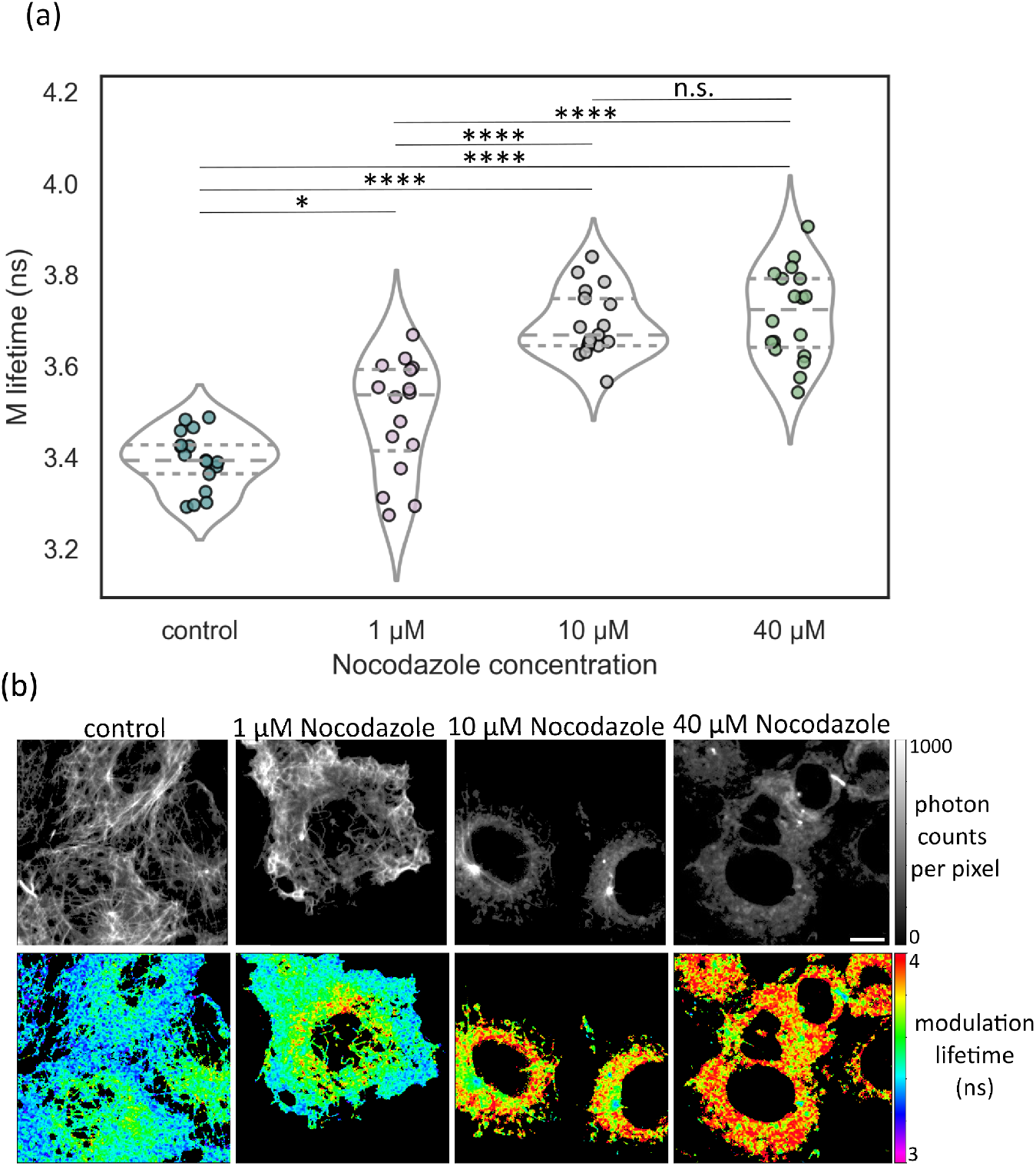
Data analysis performed on exported FLIMPA code reveals that 10 *µ*M Nocodazole is enough to completely destabilise microtubules. (a) Violin plots of SiR-tubulin modulation lifetime in COS-7 cells treated with 0 *µ*M (control), 1 *µ*M, 10 *µ*M, and 40 *µ*M Nocodazole. The plots were generated in Python using fluorescence lifetime values exported from FLIMPA. The lines within the plots indicate the interquartile range and median. The mean SiR-tubulin modulation lifetime is 3.39 ns, 3.49 ns, 3.69 ns and 3.71 ns for the control, 1 *µ*M, 10 *µ*M, and 40 *µ*M Nocodazole. The statistical analysis was performed using one-way ANOVA with Tukey multiple comparisons, where * indicates a p-value *<* 0.05, **** indicates a p-value *<* 0.0001, and n.s. denotes non-significant results. The plot was edited in Inkscape to highlight significance levels. Data were collected from four independent repeats. (b) Example intensity and modulation lifetime maps of COS-7 microtubules treated with different Nocodazole concentrations generated using FLIMPA. Scale bar is 10 *µ*m.

Finally, to validate that Nocodazole itself, regardless of microtubule depolymerisation, does not affect the lifetime of SiR-tubulin, we imaged SiR-tubulin solution on a coverslip, and no change in the probe’s fluorescence lifetime was observed when the solution was mixed with 1 *µ*M, 10 *µ*M, and 40 *µ*M of Nocodazole (Supplementary Figures 4 and 5). Nonetheless, as shown in Figure 4, SiR-tubulin aggregation was observed, providing further evidence that phasor points of lower fluorescence lifetimes are associated with clustered dye molecules rather than cellular structures.

## 3 Discussion

We have introduced FLIMPA, an easy-to-use, open-source, and standalone software package for phasor plot analysis, developed in Python. Our results demonstrate that FLIMPA is a valuable tool for researchers to investigate local variations within individual FLIM images as well as to examine fluorescence lifetime differences across multiple samples. In particular, the ROI selection tool can facilitate the identification of localised treatment effects and outliers. Additionally, the software’s advanced phasor and image visualisation options, along with the capability to present data using violin plots and tables of fluorescence lifetimes, enable a thorough comparison of the effects of experimental conditions on the fluorescence lifetimes. Finally, FLIMPA also allows users to export all generated data for further statistical analysis.

The tools provided in FLIMPA extend the capabilities of FLUTE [28], another opensource phasor plot application. Unlike the Napari-Live-FLIM plug-in [29], FLIMPA is also a stand-alone application that is straightforward to execute without requiring any coding expertise. Furthermore, and in contrast to proprietary commercial alternatives, we have ensured our application is accessible to researchers across various laboratories by supporting the analysis of multiple file formats, including .std, .ptu, and .tiff and by making the user interface intuitive and straightforward to use.

Utilising FLIMPA, we introduced a novel FLIM-based assay for quantifying microtubule network disruption upon exposure to the microtubule depolymerising agent Nocodazole. Specifically, the fluorescence lifetime of the small molecule dye SiR-tubulin was employed as a sensor of microtubule depolymerisation. Our results demonstrate that the SiR-tubulin fluorescence lifetime increases with Nocodazole concentration. In future work we will extend the assay to other well-characterised depolymerising agents, such as colchicine and vinblastine, thus we hope that FLIMPA will prove useful in the study of structural changes induced by drugs and for the search of drug candidates relevant for cancer research.

## 4 Methods

### 4.1 Phasor Plot Theory

In time-domain FLIM, the output data sets are three-dimensional, where the x, y-dimensions correspond to the spatial information while the z-dimension is the change of fluorescence intensity with time, I(t), given by Eq. (1) [2]:

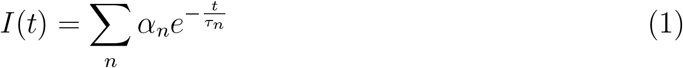

Where *t* is time, *α*_*n*_ the fractional contribution of each lifetime component *τ*_*n*_, and *n* is the number of components.

The fluorescence decay curve can be transformed to the frequency domain using the Fourier transform as expressed by Eq. (2) [40]:

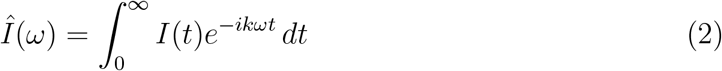

Where *ω* is the angular frequency, *i* is the imaginary component, and *k* is the harmonic number [40]. The angular frequency is derived via the repetition rate of the laser *f* as *ω* = 2*πf*. In this work, the phasor plot analysis is performed using the first harmonic, therefore *k* is set to 1.

The phasor coordinates, *g*− and *s*− of the real (cosine) and imaginary (sine) parts of the Fourier Transform for a given pixel (*x, y*) are calculated as follows by Eq. (3) and Eq. (4), respectively [41]:

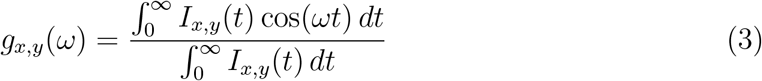

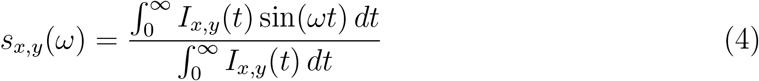

In frequency domain FLIM, the *g*− and *s*− coordinates at each pixel (*x, y*) can be calculated from the modulation ratio *m*_(*x,y*)_ and phase delay *ϕ*_(*x,y*)_ using Eq. (5) and Eq. (6) [41]:

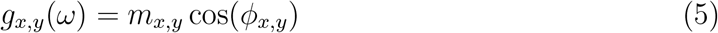

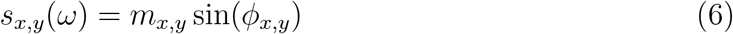

By plotting *g*− and *s*− coordinates, each pixel is represented by a point within the phasor plot [19]. The semi-circle is known as the universal circle, where points corresponding to shorter lifetimes are placed closer to the right-hand side of the universal circle (g, s = 1,0) while points of longer lifetimes are closer to the origin of the circle (g, s = 0,0), as shown in Figure 8. Moreover, phasor points of mono-exponential decays lie on the universal circle while points calculated from multi-exponential decays fall within the circle [41].

**Figure 8:**
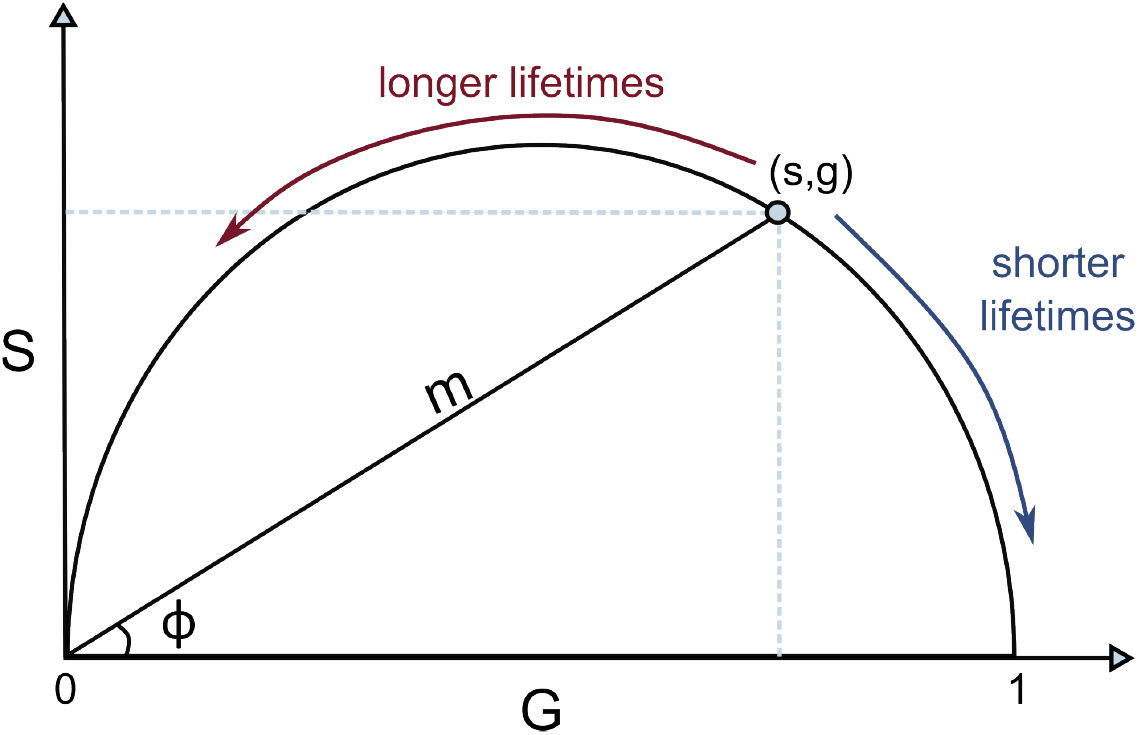
Schematic of a phasor plot, where the phase angle is denoted by *ϕ* and the modulation *m* is shown as a vector. The figure was recreated in Inkspace from Sun *et al*. 2014 [41].

Using trigonometric functions, the modulation and phase can be derived from the *g*− and *s*− coordinates of a given point by Eq. (7) and Eq. (8) [28]:

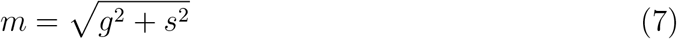

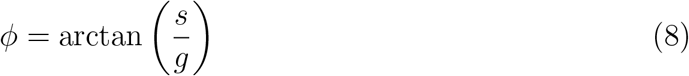

The fluorescence lifetime corresponding to the modulation and phase, given by Eq. (9) and Eq. (10) [28], respectively, are referred to as “modulation lifetime” and “phase lifetime”.

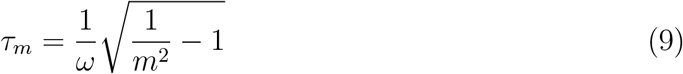

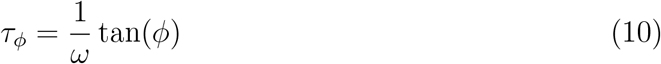

For mono-exponential species the phase and modulation lifetimes are equal [42]. Additionally, an average of the phase and modulation lifetime can be taken, which is referred to as the “average lifetime” in FLIMPA. To account for instrumental factors, such as differences in the detector and optics, a reference file with a known lifetime is used for calibration. Commonly, Rhodamine 6G which has a known lifetime of 4 ns is used as the reference sample. Other unquenched dyes can also be used, with the reference correction serving a similar purpose as the Instrumental Response Function (IRF) deconvolution in curve-fitting methods.

### 4.2 Implementation

FLIMPA was developed using Python (version 3.11.7), with the phasor plot analysis predominantly performed using the NumPy [43] (version 2.0.0) and SciPy [44] (version 1.14.0) libraries. The frontend of the application was built using PySide6 (version 6.7.2), while plotting and visualisation were achieved using Matplotlib [45] (version 3.9.1), Seaborn (version 0.13.2) [46], and Pandas [47] (version 2.2.2). The complete list of libraries used for building FLIMPA is available in the requirements.txt file on GitHub. Statistical analysis of the exported data was conducted using the Pingouin library [48] (version 0.5.4).

### 4.3 Manual mask generation

To remove areas of SiR-tubulin molecule clustering, manual masks of the COS-7 microtubules were created. This involved first segmenting the background of the microtubules in Python (version 3.11.7) by setting a threshold of 200 photon counts per pixel, followed by importing the masks generated to FLIMfit [20] and manually removing the areas that corresponded to the dye aggregation.

### 4.4 System requirements

FLIMPA can be executed on a Windows computer using the .exe file provided on its GitHub repository. The repository also includes FLIMPA’s backend and frontend Python code. The sample .sdt files used for Figures 1, 2, 3, and 5 are provided so that the GUI can be easily tested. The user manual for FLIMPA can be found in the Supplementary information section. The processing speed of FLIMPA was evaluated using 72 Becker & Hickl .sdt files with dimensions of 256×256×256, where each image took an average of 1.4 seconds to process without pixel binning, and 1.8 seconds with 3×3 pixel binning applied. These results were obtained using a system running Windows 10, equipped with an Intel(R) Core(TM) i7-9750H CPU @ 2.60GHz.

### 4.5 Cell culture

COS-7 cells were acquired from the American Type Culture Collection (ATCC, USA) and cultured at 37 °C with a 5% CO_2_ supply. Cells were grown in Dulbecco’s modified Eagle’s medium (DMEM, ThermoFisher Scientific, USA) supplemented with 10% fetal bovine serum (FBS, Thermo Fisher Scientific), 1% penicillin-streptomycin and 1% of GlutaMAX™ (ThermoFisher Scientific). 10K cells were seeded in each chamber of a 8-chamber glass well plate (IBIDI GmbH, Germany) with 200 µL of cell media and incubated at 37 °C with 5% CO_2_ for 24 hours. This was followed by overnight staining with 1 µM SiR-tubulin (Spirochrome, Switzerland) and 10 µM Verapamil (Spirochrome) in cell media. Verapamil is an efflux pump inhibitor and was added to the staining solution to prevent the cells from expelling the fluorescent probe. The following day, cells were washed twice with phosphate-buffered saline (PBS, ThermoFisher Scientific) and 200 µL of fresh cell media with 10 µM Verapamil was added. After media replacement, the cells were treated with 1 *µ*M, 10 *µ*M, or 40 *µ*M of Nocodazole for 30 minutes. Imaging occurred immediately following treatments, without removal of the Nocodazole, during which an on-stage incubator system (OKOLab, Italy) was used to maintain the cells at 37 °C with 5% CO_2_ supply.

### 4.6 In-house TCSPC set-up

The in-house TCSPC module is based on a confocal microscope. The samples were illuminated with a pulsed laser (Fianium Whitelase, Denmark) at 40 MHz frequency [49]. Imaging was performed using an Olympus IX83 microscope system with a 60x oil objective with a 1.40 numerical aperture (Olympus, Japan). The fluorescent photons were captured using a Becker & Hickl GmbH PMC-150 photon multiplier tube (PMT) and SPC-830 Photon Counting Electronics [49]. Photons are accumulated over 10 cycles with a duration of 12 seconds each, with output data having the dimensions of 256×256×256 (t, x, y). The excitation and emission filters were centred at 635 nm and 700 nm (FF02-632/22-25, Semrock Inc, USA and ET700/75m, Chroma, USA). [49]. Rhodamine 6G (at a concentration of 10^5^ *µ*M in H_2_O) was used as the reference sample and measured using excitation and emission filters centred at 510-20 nm and 542-27 nm (FF03-510/20-25 and FF01-542/27-25, Semrock Inc), respectively. The image of the Convallaria Rhizome sample (shown in Supplementary Figure 2) was acquired using the 510-20 nm and 542-27 nm filters and a 40x oil Olympus objective with a 1.40 numerical aperture. To prevent photon pile-up, the photon counts at the detector were kept below 1% of the instrumental frequency.

## Supporting information

Supplementary Figure 1, Supplementary Figure 2, Supplementary Figure 3, Supplementary Figure 4, Supplementary Figure 5, user manual

## 6 Supplementary information

Supplementary figures and user manual.

## 7 Data availability

The executable file, Python code and sample data can be found on FLIMPA’s GitHub repository.

## 8 Acknowledgements

G.S.K.S. acknowledges funding from the Wellcome Trust (065807/Z/01/Z) (203249/Z/16/Z), the UK Medical Research Council (MRC) (MR/K02292X/1), Alzheimer Research UK (ARUK) (ARUK-PG013-14), Michael J Fox Foundation (16238 and 022159), and Infinitus China Ltd. C.F.K. acknowledges funding from the UK Engineering and Physical Sciences Research Council (EP/L015889/1 and EP/H018301/1), the Wellcome Trust (3-3249/Z/16/Z and 089703/Z/09/Z), the UK Medical Research Council (MR/K015850/1 and MR/K02292X/1), and Infinitus China Ltd. S.K. acknowledges funding from AstraZeneca and the UK Engineering and Physical Sciences Research Council (EPSRC) grant EP/S023046/1 for the Centre for Doctoral Training in Sensor Technologies for a Healthy and Sustainable Future. N.F.L. acknowledges the Swiss National Science Foundation (Grant Number P2EZP2 199843).

