## Supplementary Figure 1, Supplementary Figure 2, Supplementary Figure 3, Supplementary Figure 4, Supplementary Figure 5, user manual for "FLIMPA: A versatile software for Fluorescence Lifetime Imaging Microscopy Phasor Analysis"

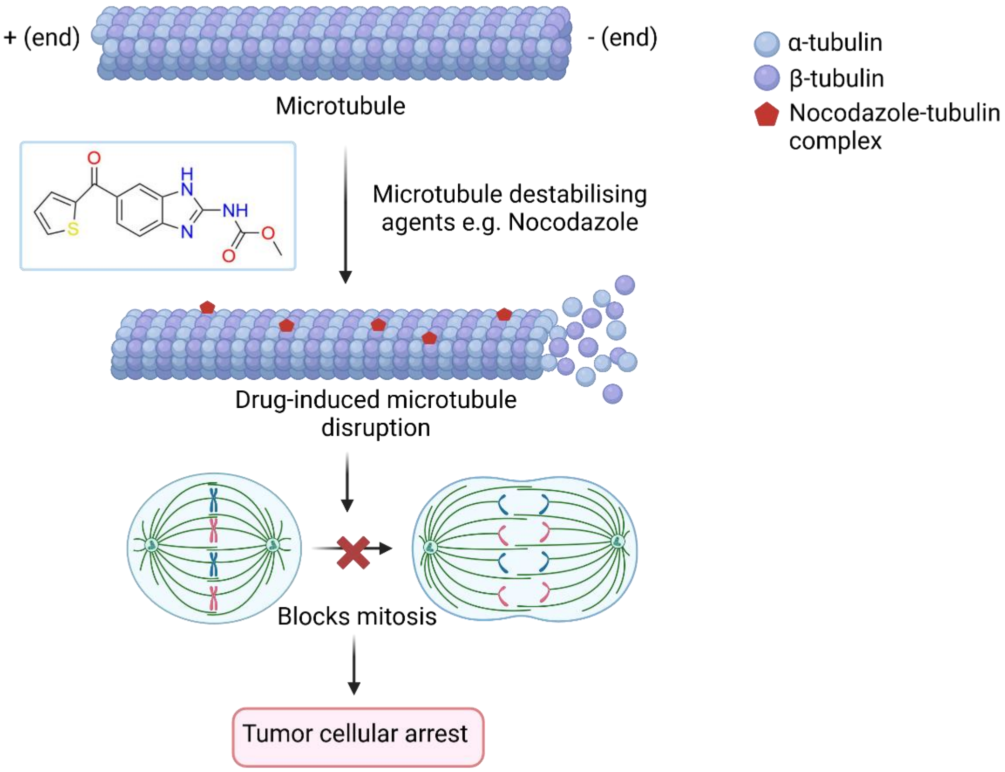

**Supplementary Figure 1** Overview of case study on quantifying microtubule depolymerisation upon drug exposure. Microtubule destabilising agents, such as Nocodazole, bind to the microtubules and induce the disassembly of the tubulin subunits leading to mitotic arrest and cell death. A FLIM-based assay was used to quantify the drug-induced microtubule depolymerisation, using the fluorescence lifetime of SiR-tubulin, a small dye molecule that selectively binds to the β-subunits of microtubules. The schematic was recreated from Gupta et al. (2019) [1] using BioRender with the Nocodazole structure designed in ChemDraw.

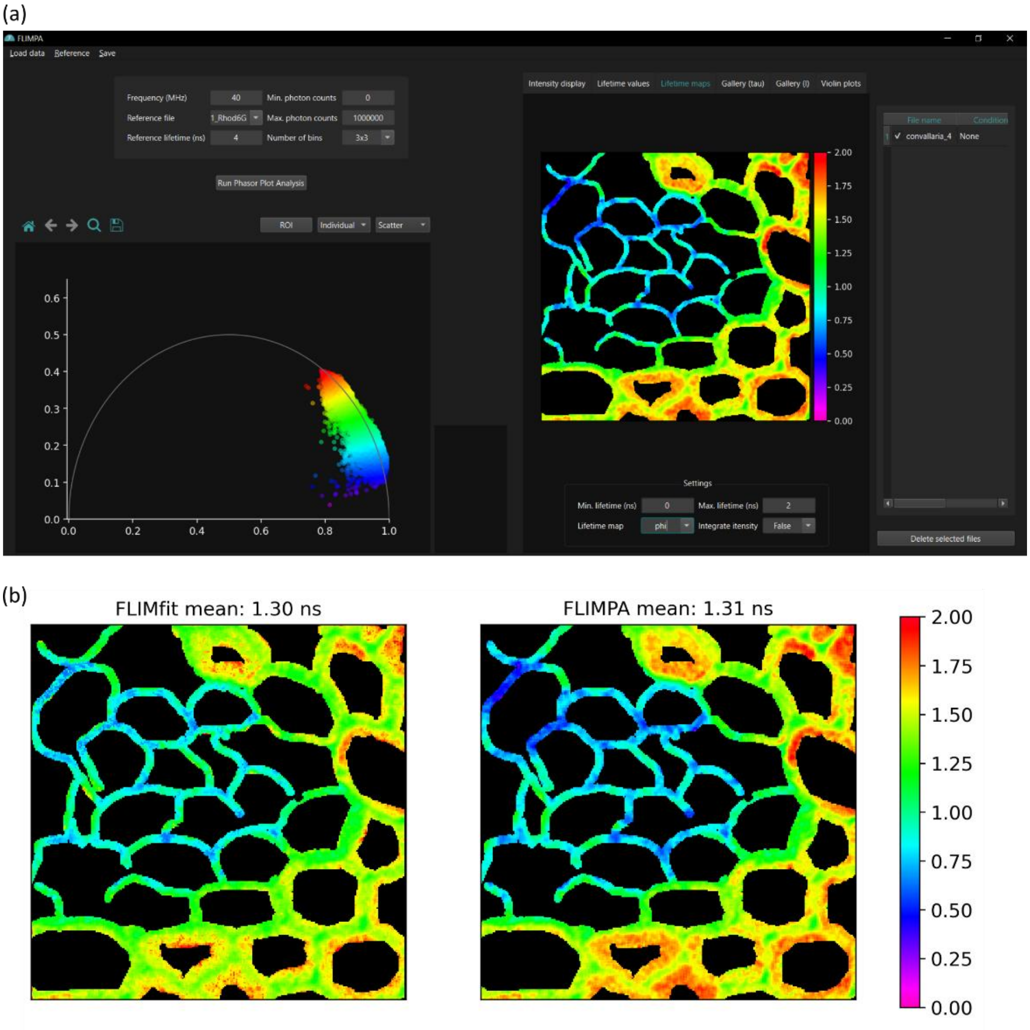

**Supplementary Figure 2** (a) Overview of the FLIMPA GUI displaying the phasor plot analysis of a Convallaria Rhizome sample. (b) Comparison between the bi-exponential fluorescence lifetime image of Convallaria Rhizome from FLIMfit [2] (left) and the phase lifetime map generated by FLIMPA (right). The mean bi-exponential fluorescence lifetime and mean phase lifetime are indicated in the image subtitles. Both images were plotted in Python, with the colour bar representing time in nanoseconds.

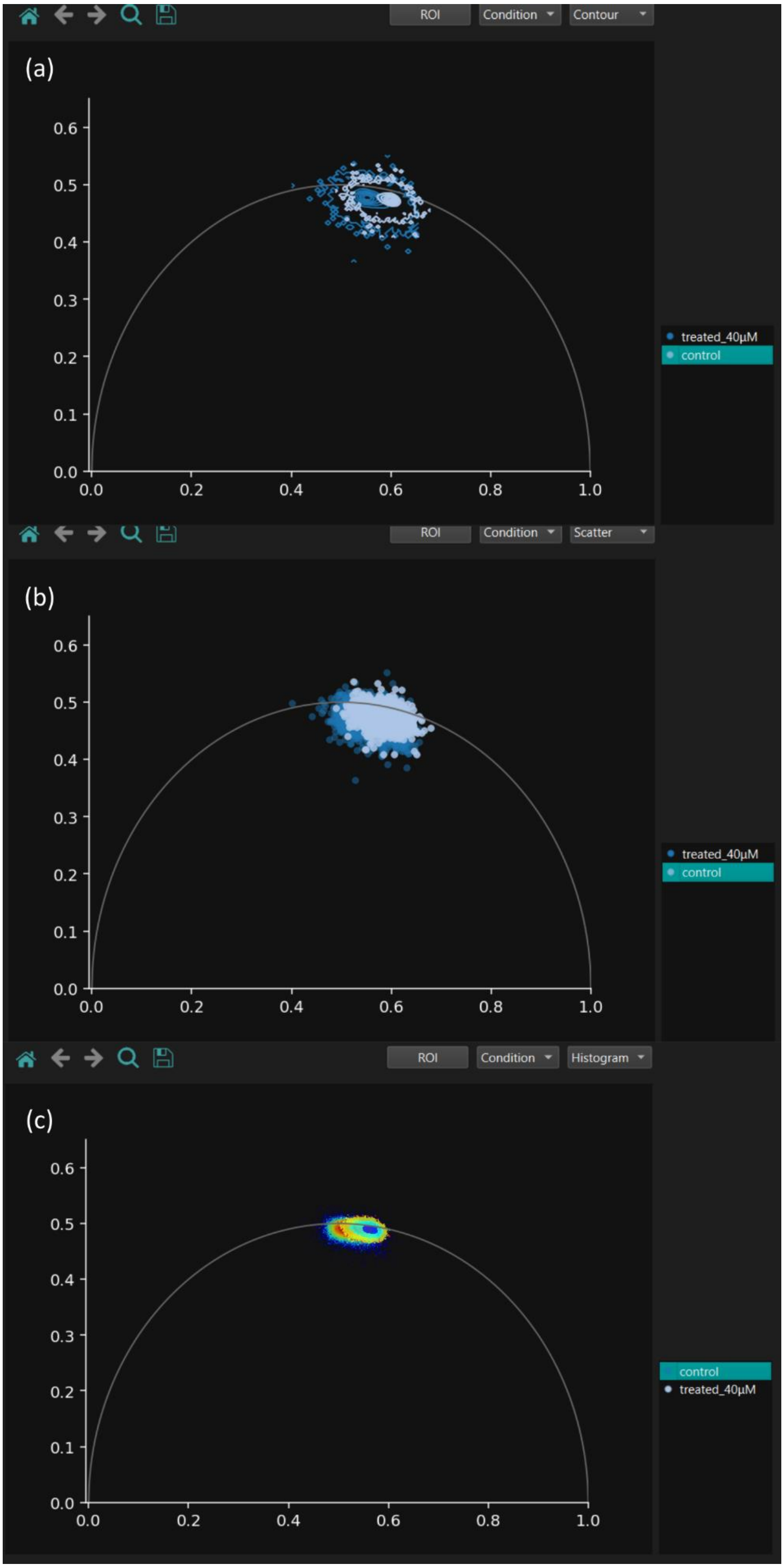

**Supplementary Figure 3** Phasor clouds visualisation options provided by FLIMPA using (a) contour maps, (b) scatter plots and (c) histogram.

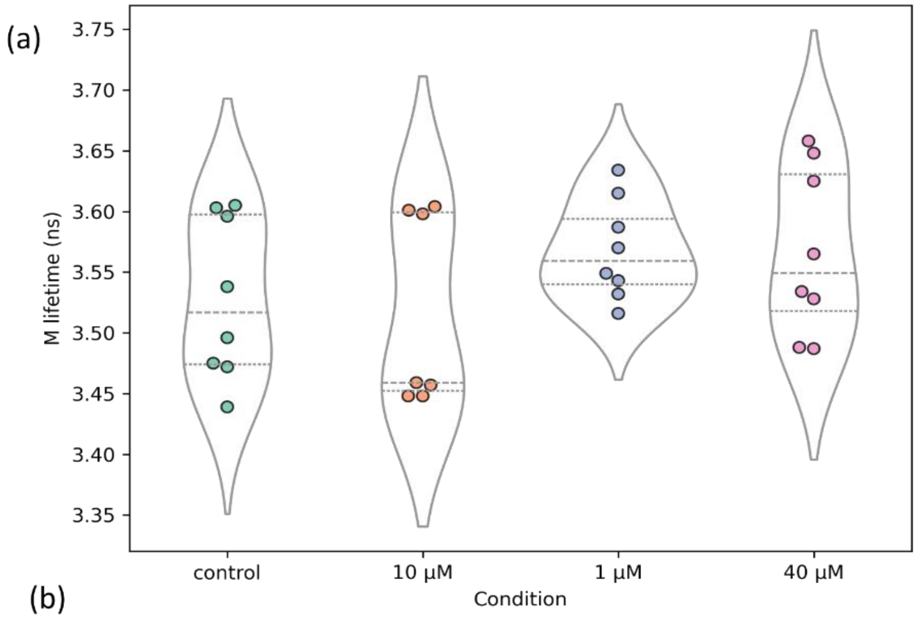

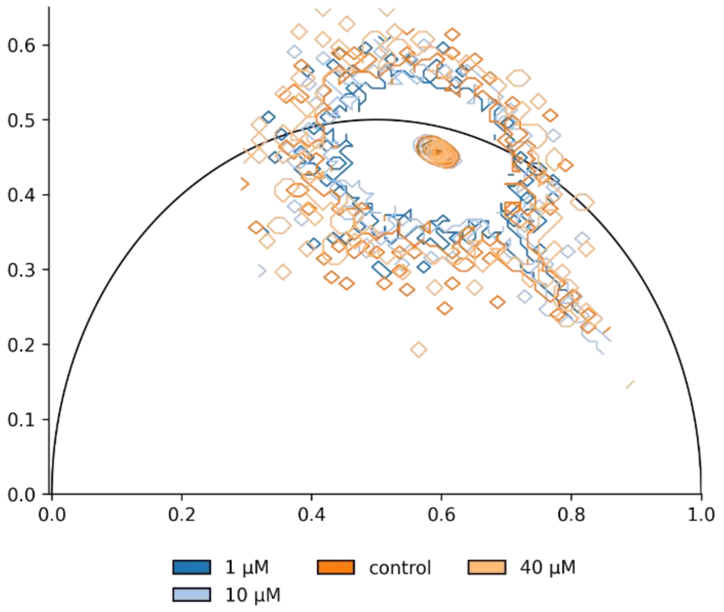

(c)

| Condition | M lifetime (ns) |
|---|---|
| control | 3.53 |
| 1 μM | 3.57 |
| 10 μM | 3.52 |
| 40 μM | 3.57 |

**Supplementary Figure 4** Studying the effect of Nocodazole on SiR-tubulin fluorescence lifetime imaged on a coverslip. (a) Violin plots showing the effect of different concentrations of Nocodazole on SiR-tubulin modulation lifetime; (b) phasor plot exported from FLIMPA for control (darker orange), 1 μM Nocodazole (darker blue), 10 μM Nocodazole (light blue), and 40 μM Nocodazole (light orange); (c) table of modulation lifetimes in nanoseconds per Nocodazole concentration.

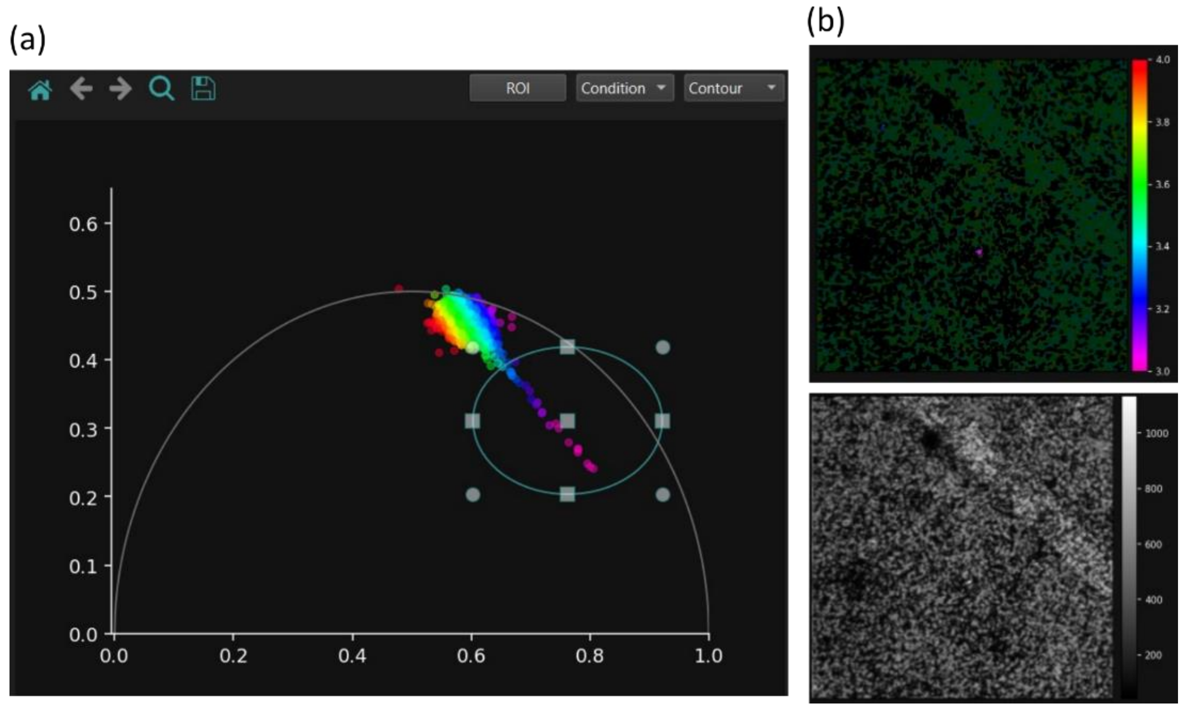

**Supplementary Figure 5** SiR-tubulin molecule aggregation. (a) Phasor plot from a sample of SiR-tubulin imaged on a coverslip where the phasor points with lower lifetime, selected using the ROI tool, correspond to dye aggregation. (b) Corresponding modulation lifetime map highlighting the pixels associated with dye clustering (top) and intensity map (bottom).

**User Manual**

Please note that an **online user manual** with in-depth animations can be accessed here.

### 1. Getting started

From FLIMPA's GitHub repository, the following material can be downloaded:

- The .exe file for FLIMPA running on Windows.
- Alternatively, FLIMPA can installed from its GitHub repository following the instructions provided there.
- Sample Becker & Hickl .sdt files alongside their .tif masks.

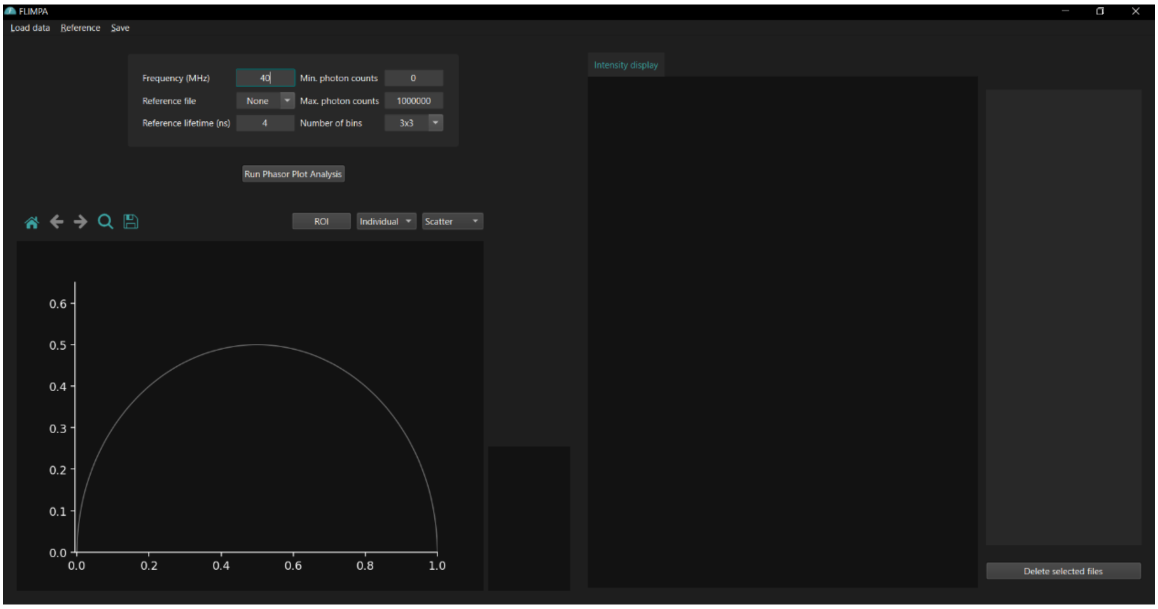

**Figure 1** FLIMPA GUI upon launching the software.

### 2. Loading data

Raw data file formats currently supported:

- **.sdt, .ptu, .tif**

While loading the data, users can choose from the options:

- Define the **experimental condition** for the raw data (e.g., 'control'), allowing images with the same condition to be visualised together later and/or
- Provide **manual masks**, which currently need to be created with FLIMFit. The manual masks must be in .tif format

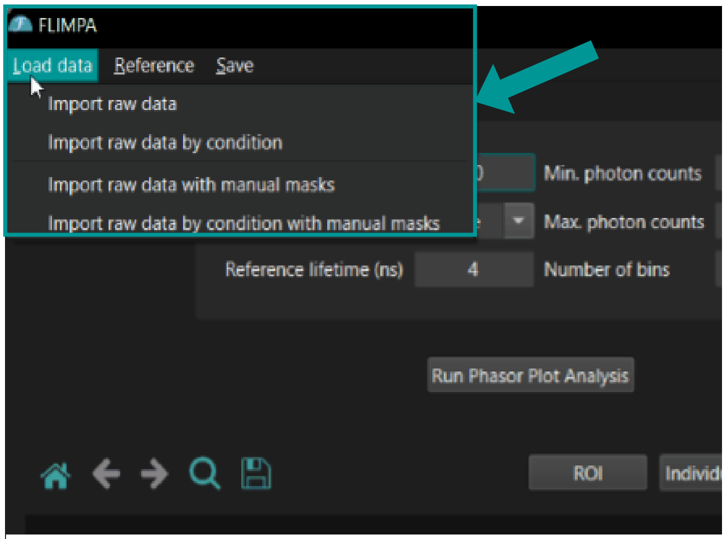

**Figure 2** Load data by clicking the 'Load Data' button located in the top left corner of the app.

### 2.1. Loading data alongside manual masks

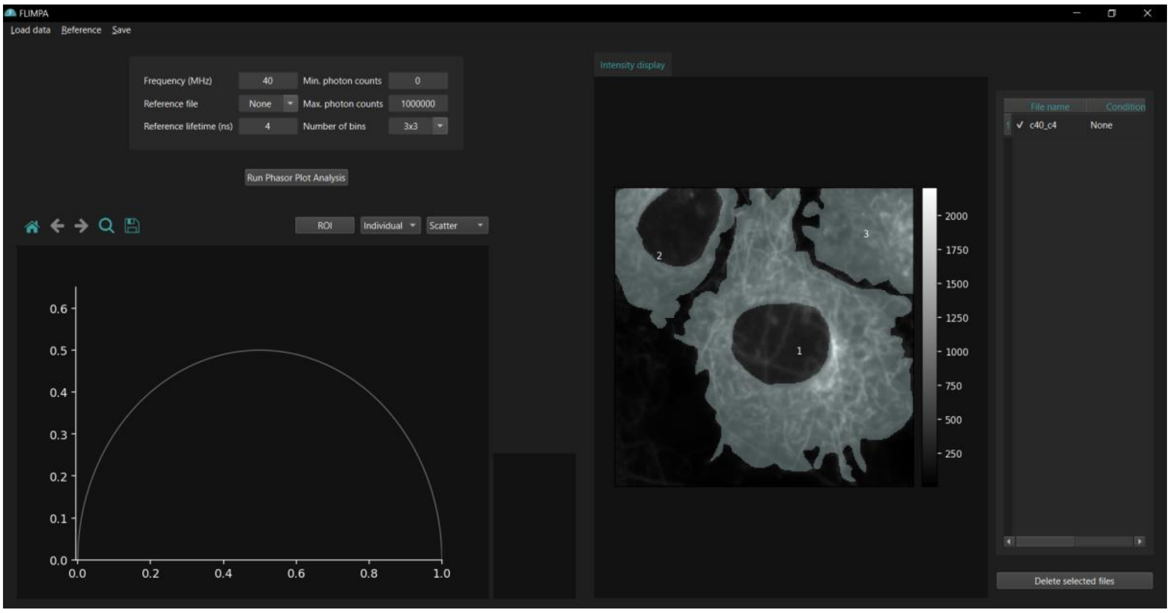

**Figure 3** Raw FLIM data can be loaded with their respective manual masks. If the manual masks consist of more than one region of interest (ROI), these will be numbered accordingly.

- Manual masks can be created using FLIMFit and should be named according to the raw data, followed by “ segmentation.tif" (e.g., "control.sdt" should have a mask named "control segmentation.tif")
- Click on “Load data” on the left corner of the GUI, followed by “Import data with manual masks”
- Select the raw data, then choose the folder containing the masks
- Once loaded, different ROIs (e.g., cells) will be numbered for separate analysis
- Alternatively, users can select "Import raw data by condition with manual masks" to assign both the experimental condition and provide the corresponding masks

### 3. Running the phasor plot analysis

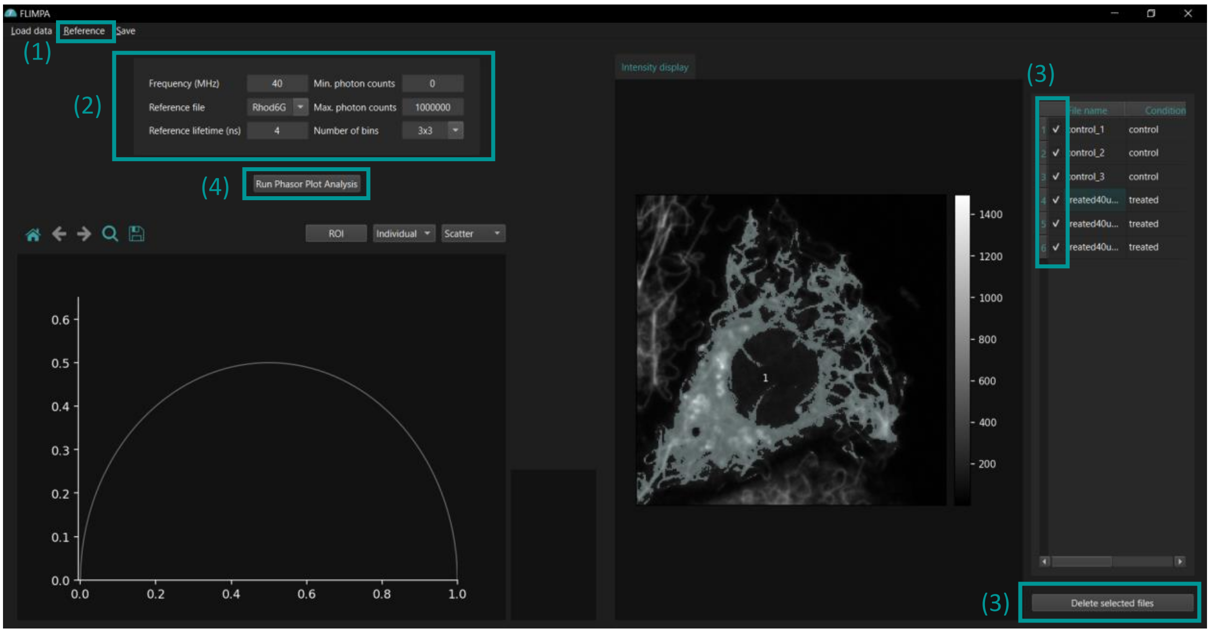

**Figure 4** Overview of steps required for running the phasor plot analysis.

1) Load a reference file, such as a dye with a known lifetime (e.g., Rhodamine 6G with a reference lifetime of 4 ns) or an IRF file. Accepted formats for IRF files are .sdt and .csv, with the reference lifetime set to 0 ns. The .csv files should contain two columns: the first column representing the time intervals at which the IRF was recorded, and the second column containing the corresponding intensity values. After loading the IRF, FLIMPA will prompt users to specify whether the data are stored in "Column 0" or "Column 1"; please select "Column 1"
2) To run the phasor plot analysis, users will need to specify the following parameters:
   - Laser Frequency (in MHz)
   - Reference File Lifetime (in ns)
   - Number of Time Bins (select from None, 3x3, 7x7, or 9x9)
   - Minimum Photon Count Threshold (optional, at least 100 p.c. per pixel recommended)
   - Maximum Photon Count Threshold (optional)
3) The check box has a dual function:
   - Only selected files will be analysed
   - Users can click on “Delete selected files” to remove files not needed
4) Click to run the phasor plot analysis

### 4. Results overview

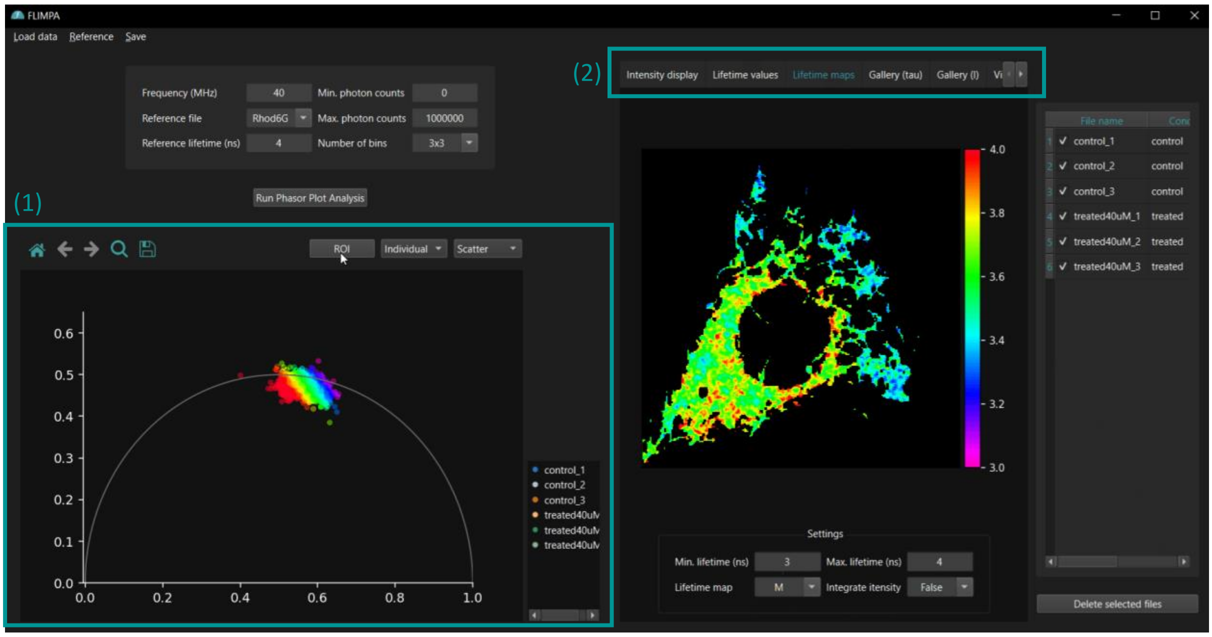

**Figure 5** Upon completion of the phasor plot analysis, section (1) will display the phasor plot visualisations, while area (2) will present newly generated tabs containing FLIMPA's analysis results

The phasor plot analysis results are displayed across different tabs, specifically the "Lifetime values" table, "Lifetime maps", "Gallery (tau)", "Gallery (I)", and "Violin plots" tabs.

#### 4.1. Fluorescence "Lifetime values" table

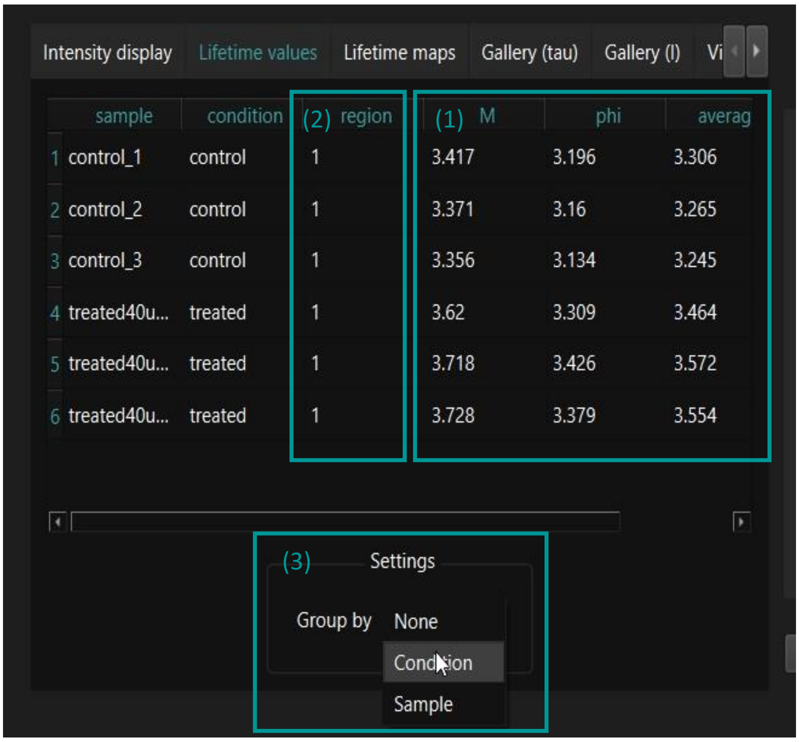

**Figure 6** Tab showing the mean fluorescence values per image or ROI in a table format.

1) The table will display the modulation (M), phase (phi), and the average of the modulation and phase fluorescence lifetimes for each image
*2)* If different ROIs were provided using manual masks (as shown in section 2.1. Loading data alongside manual masks), the mean fluorescence lifetimes will be listed per ROI instead of per image
3) Additionally, users can select "Group by Condition" to display the mean lifetime per experimental condition or "Group by Sample" to calculate the mean lifetime per image if different ROIs per image have been provided

#### 4.2. Fluorescence "Lifetime maps"

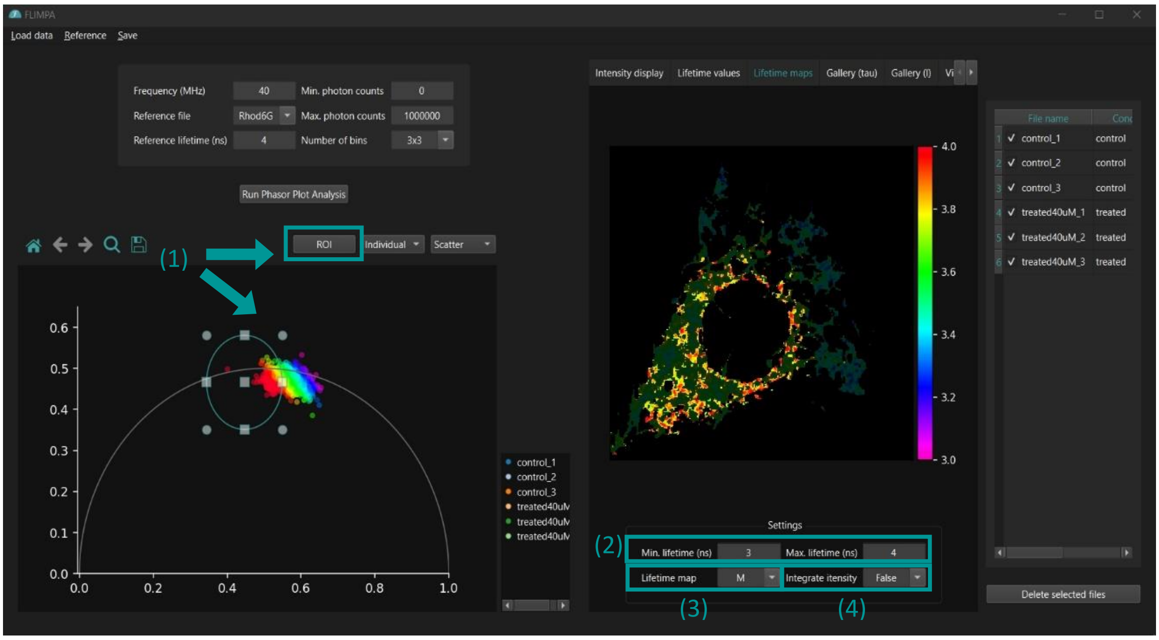

**Figure 7** ''Lifetime maps" tab displays the the individual images where localised effects can be explored using the ROI selection tool.

1) ROI selection tool can be used to investigate localised effects within individual images
2) Users can set the minimum and maximum lifetime range (in ns) for the fluorescence lifetime map colour bar
3) Users can select between visualising the modulation (M), phase (phi) or average of modulation and phase lifetime map
4) If "integrate intensity" is set to "True", the fluorescence lifetime map will be integrated with the intensity image

### 4.3. Gallery (tau) tab

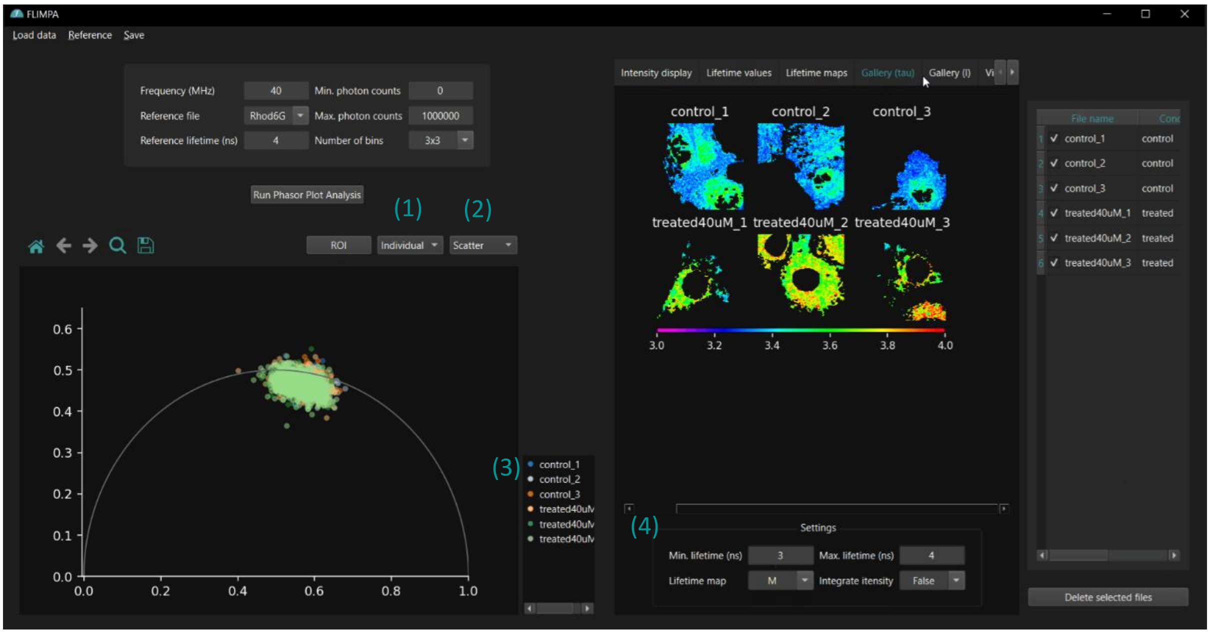

**Figure 8** Overview of the "Gallery (tau)" tab, which contains the gallery of the fluorescence lifetime maps of the different samples analysed. The phasor clouds of the samples are displayed in a single plot and can be coloured based on their sample identities (as shown in this Figure) or based on their experimental condition.

1) Phasor clouds can be coloured based on their sample identity (select "Individual") or by their experimental condition (select "Condition")
2) Phasor visualisation options are scatter plots, histograms or contour maps
3) Legend to highlight entry selected
4) The gallery of lifetime maps has the same user setting options as the individual lifetime maps shown in Figure 7

#### 4.4. Gallery (I) tab

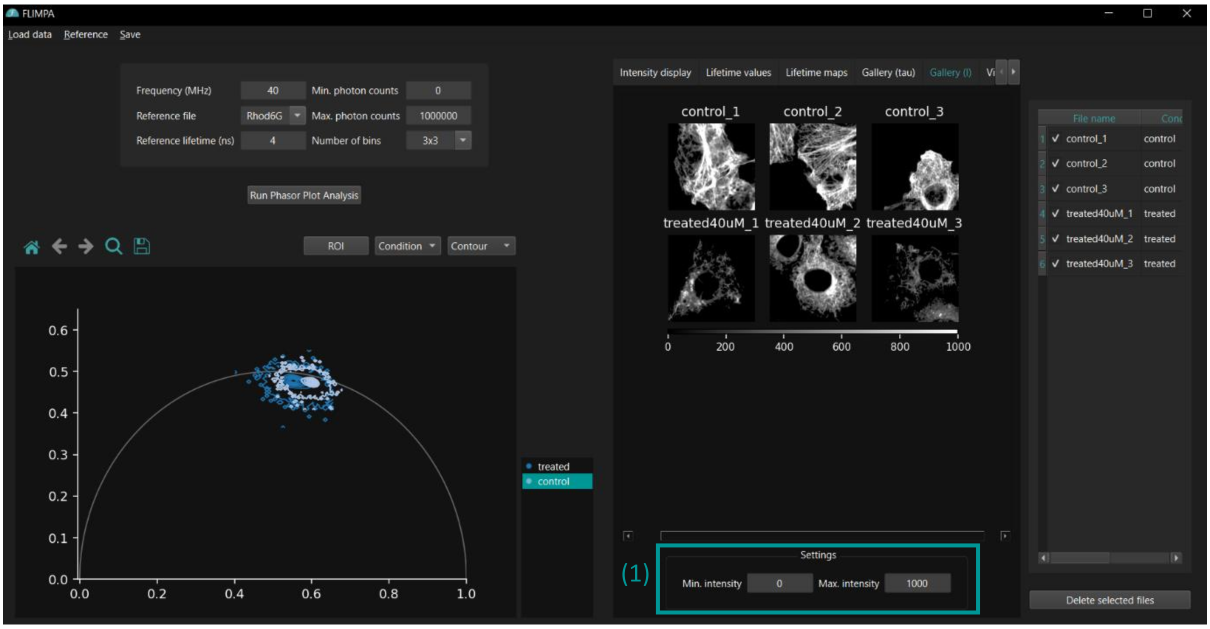

**Figure 9** "Gallery (I)" tab provided a gallery of fluorescence intensity maps.

1) Users can set minimum and maximum photon counts per pixel for the fluorescence intensity image colour bar

#### 4.5. Violin plots

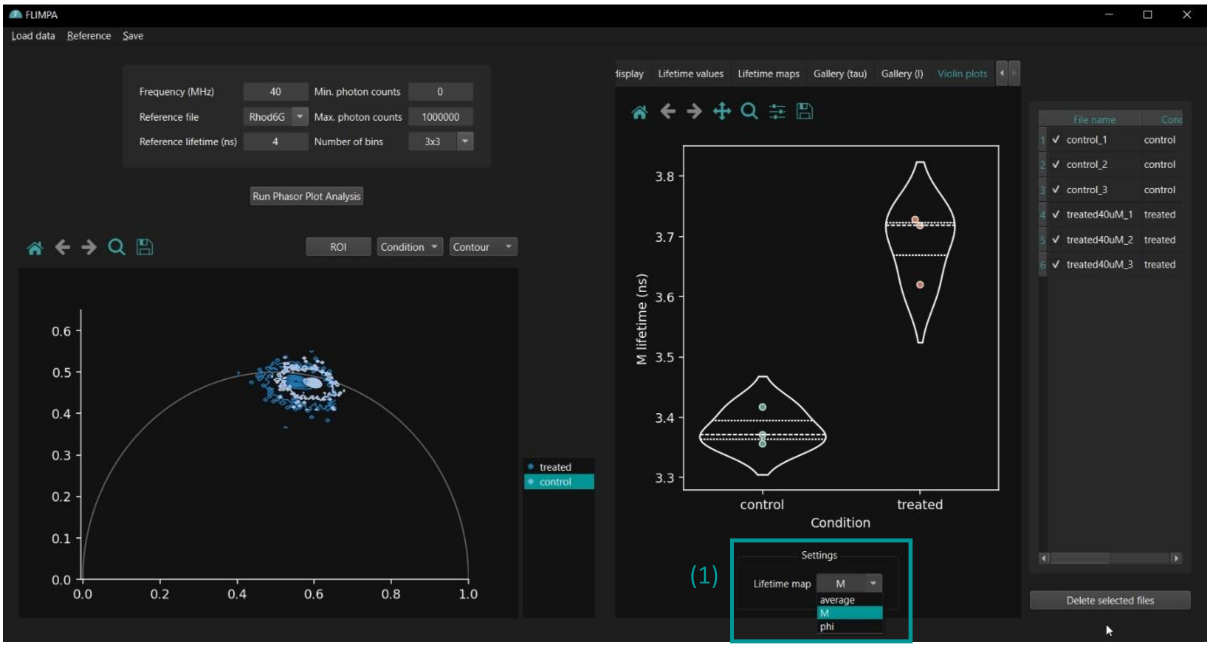

**Figure 10** The "Violin plots" tab displays the distribution of data points for each treatment group.

1) Users can select between plotting the mean average, modulation (M) or phase (phi) lifetime per image, or per ROI if manual masks with multiple ROIs have been provided

### 5. Saving FLIMPA's outputs

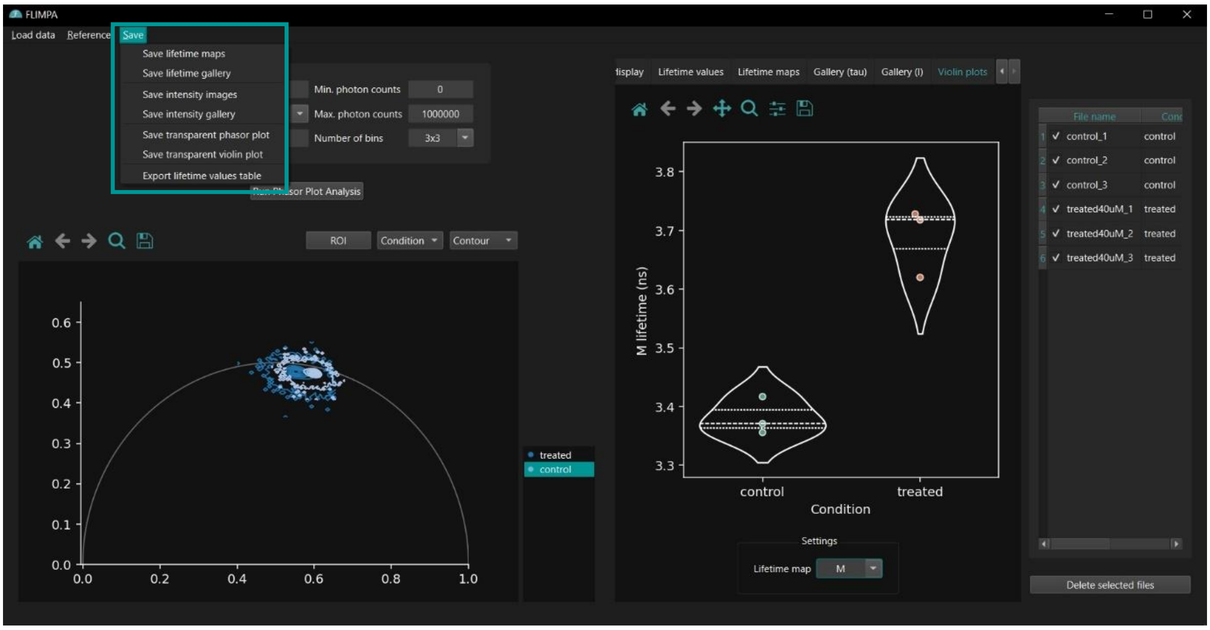

**Figure 11** The data generated can be saved by clicking on the "Save" button on FLIMPA's toolbar.

FLIMPA allows users to export all generated data. This includes:

- Lifetime and Intensity Maps: Exported as .png and raw .tif files
- Gallery Visualisations: Lifetime and intensity galleries exported as .png files
- Phasor Plots and Violin Plots: Saved with a transparent background
- Statistical Data: A .csv file containing the mean fluorescence lifetime per image can be exported for further statistical analysis
